# Proliferation to suppress neoplasia: a general model and a first test in the moon jelly

**DOI:** 10.1101/2025.09.07.674745

**Authors:** Anish A. Sarma, Lea Goentoro, John C. Doyle

## Abstract

Cell proliferation is necessary to maintain tissue homeostasis, but proliferation carries with it a risk of cancer. Nevertheless, nature abounds with examples of organisms that achieve low rates of cancer while also proliferating to maintain tissue homeostasis. To understand how organisms might achieve both of these goals, we developed a dynamical model describing cell birth, death, and mutation in a population. The model identifies two distinct regimes. In one regime, as expected, decreasing proliferation delays accumulation of neoplastic cells. In another regime, unexpectedly, increasing proliferation suppresses accumulation of neoplastic cells. In this regime, when more cells proliferate, more cells correspondingly die as a consequence of homeostatic feedback. As long as neoplastic cells are detected and killed preferentially, the high flux of cells acts as a proofreader, eliminating neoplastic cells. **High-flux proofreading** may seem costly, but it can be effective even when the system does not have a precise detector of neoplastic cells. As a first experimental test of whether high-flux proofreading is biologically relevant, we examined the moon jelly, a cnidarian. Neoplasms have rarely been observed in cnidarians, and yet simply inhibiting proliferation is sufficient to promote neoplasms in the moon jelly. Together, the model and experiments show that high-flux proofreading is an effective cancer resistance strategy. Because cell birth, death, and mutation are fundamentally conserved processes, high-flux proofreading may be widespread. The quantitative framework presented in this study offers clear experimentally testable predictions to assess high-flux proofreading in other systems, and its potential utility for cancer prevention and treatment.

## Introduction

Cells in tissues experience continuous wear and tear in the course of typical life. Maintenance of tissue structure and function requires cells to be continually replaced.^1,2^ Yet cell proliferation carries risks. When a cell divides, its daughter cells can acquire or propagate mutations that, as they accumulate, can drive the development of cancer. In humans, many common cancers originate in proliferative tissues, such as the large intestine, the lungs, and the skin, while rarely originating from non-proliferative tissues, such as central nervous system neurons.^3–6^ Thus, lifetime accumulation of cell divisions appears to correlate with lifetime cancer risk.

However, it is increasingly appreciated that the relationship between proliferation and cancer may turn out to be more complex.^7–10^ In humans, there are notable exceptions to the rule, such as small intestinal epithelial tissue, in which cancers are relatively rare despite high proliferation.^3,11^ Cancer rates also vary widely across species.^12,13^ For example, elephants, lobsters, and naked mole rats are long-lived and have an unusually low rate of cancers.^8,12–15^ Jellyfish are another intriguing example: they are long-lived and capable of dramatically ramping up proliferation at opportune times, yet cancer is rarely observed at all.^16–21^

To better understand the relationship between the need to renew cells and the risk of cancer, we developed a mathematical framework to analyze general conditions that suppress cancer while also maintaining tissue homeostasis. Mathematical models have often been used to study the general population dynamics of mutation accumulation.^22,23^ Models of cancer evolution likewise begin with a population of cells, which can proliferate, die, or acquire mutations. Such models have been used to study the relationship between mutation rates and cancer prevalence,^24–26^ how cancers grow and evolve,^27^ and how cancers respond to treatment.^28–30^ Our model diverges from existing approaches in two ways. First, we implement tissue homeostasis explicitly, with feedback coupling between the rates of cell proliferation and cell death. Second, instead of asking, under what conditions do the population dynamics lead to the development of cancer, we ask, under what conditions do the population dynamics suppress the development of cancer?

## Results

### A dynamical model of birth, death, and mutation processes

Consider a population of cells consisting of healthy cells *x_0_* and neoplastic cells *x_1_* (Figure 1). Cells in each subpopulation *x_i_* undergo basic turnover processes: they divide and die, at rates represented by the functions ***a_i_***and ***w_i_***respectively. Initially all cells are healthy, but healthy cells mutate at a rate represented by the function ***m***. This birth-death-mutation model is described by the following differential equations:

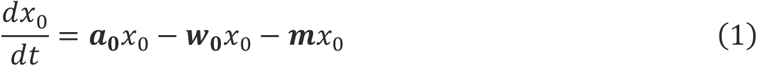

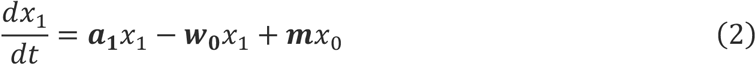

**Figure 1.**
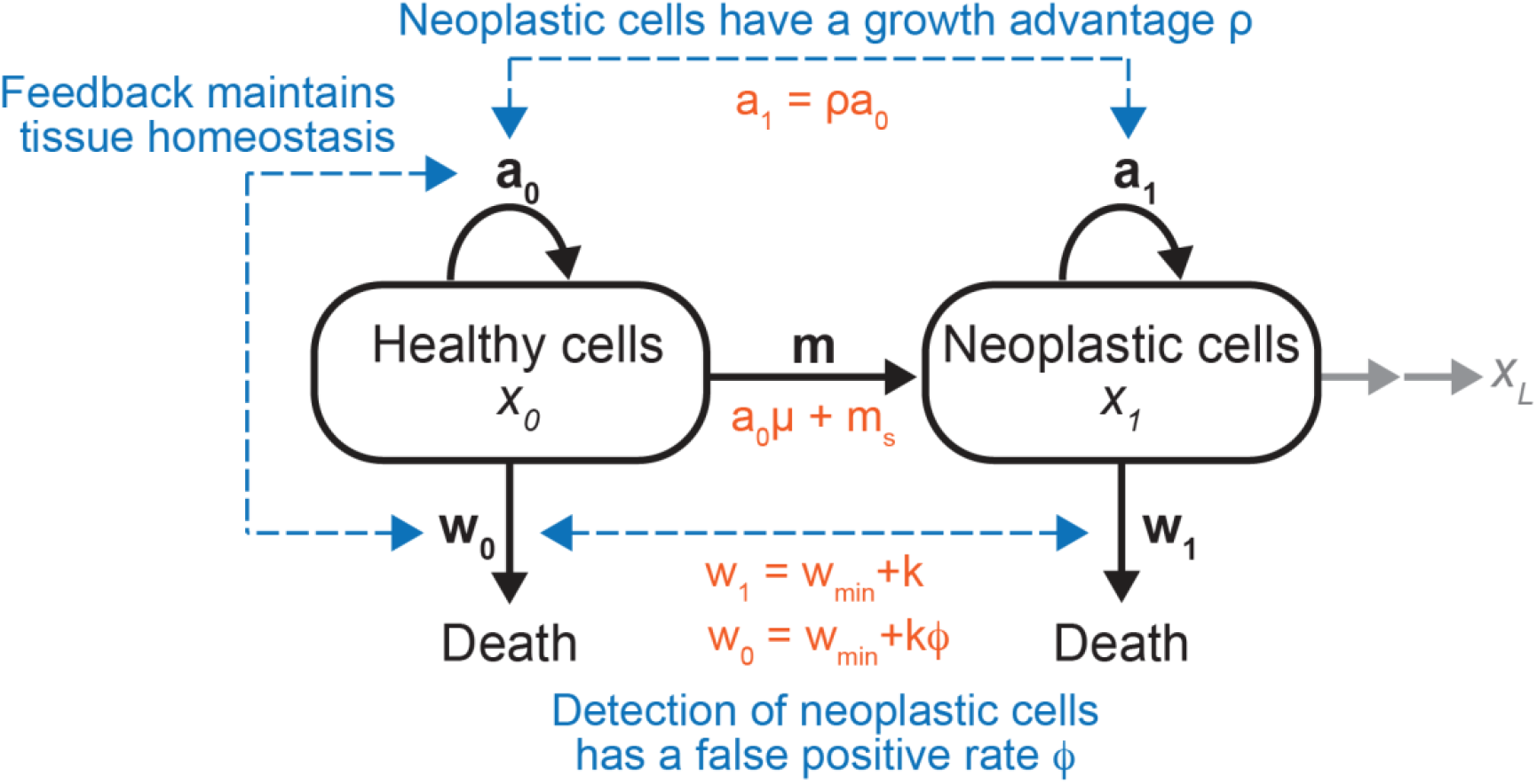
The mathematical model. In this study, we developed a mathematical model of birth, death, and mutation to explore the conditions that suppress accumulation of neoplastic cells. Each process is governed by a rate function denoted in black, bold characters. Blue lines represent coupling between the rate functions. Orange functions represent the rate functions and the linear coupling functions that we used to derive the solutions presented in the main text. See main text for more detailed description of the model.

Here, ***a, w***, and ***m***are rate functions, coupled to one another as defined next. Equations 1–2 are written with one mutation step, but the results readily extend to a model with multiple, sequential mutation steps (*x_0_, x_1_, x_2,_* … *x_L_;* Supplemental Section D). Equations 1–2 also assume that all cells can divide, but division is typically limited to stem cell populations; we verified that the key conclusions hold for a model with separate dividing and differentiated subpopulations (Supplemental Section F).

#### Rate functions

Next, we define the rate functions and the couplings between them. To facilitate the analytical discussion, we present here the modeling results with linear coupling functions. We verified that the key conclusions hold even when the model is generalized to nonlinear coupling functions (Supplemental Section E).

We define the division rate of neoplastic cells ***a_1_***as proportional to the division rate of healthy cells ***a_0_***, with a proportionality constant *ρ*. To simplify the notation, we define ***a****=* ***a_0_***.

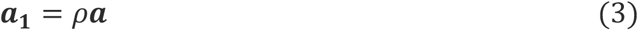

When *ρ > 1*, neoplastic cells have a growth advantage relative to healthy cells, while *ρ < 1* indicates a growth disadvantage.

For the mutation function ***m***, we take into account that mutation can occur during cell division because of replication error, described by the per-division rate *μ*, or spontaneously from exposure to environmental factors, such as carcinogens, pollutants, or pathogens, described by the rate constant *m_s_*.

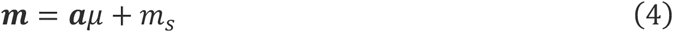

We define the death rate functions with two components, the constitutive death *w_min_* and the targeted death ***k***.

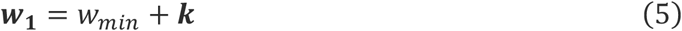

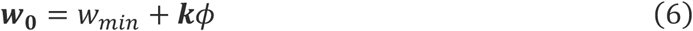

Neoplastic cells can be detected and targeted for death, for instance, by DNA damage detection mechanisms, which can trigger apoptosis.^31^ Since detection is rarely perfect, targeting neoplastic cells can inadvertently lead to some healthy cells being killed as well. Accordingly, in Equation 6, we include in the death rate function of healthy cells a false-positive rate of detection, 0 < *ϕ* < 1. Lower values of *ϕ* represent more successful discrimination between healthy and neoplastic cells.

#### Tissue homeostasis

Lastly, we implement into the model the constraint of tissue homeostasis. Tissue homeostasis can be used broadly to describe processes that maintain a tissue’s structure and function. In this model, we focus on maintenance of cell population size. To implement tissue homeostatic feedback, we first scale the population size to 1. Tissue homeostasis is achieved when the population size is stable:

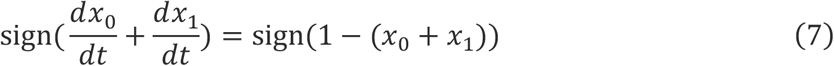

To maintain tissue homeostasis in the model, we implement into Equations 1–2 the feedback functions ***a_th_, w_th0_***, and ***w_th1_***as follows:

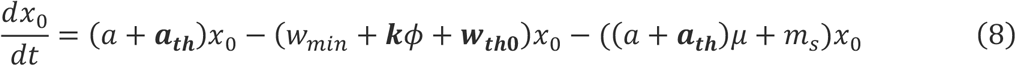

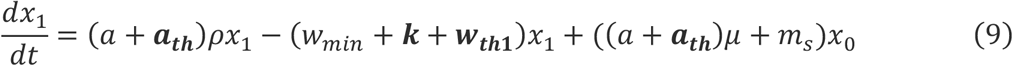

Note the convenience of defining the feedback functions in this way: when *x_0_ + x_1_ = 1*, we treat *a* as a tunable parameter (reflected by the un-bolded notation), with the remaining *x-*dependent proliferation accounted for by ***a_th_***.

Multiple feedback mechanisms have been identified that mediate tight coupling between cell proliferation and cell death, in systems as varied as the fly midgut, the hydra, and human embryonic stem cells.^32–34^ Because there are varied ways biological systems implement this feedback, for simplicity, we use the following generic functions.

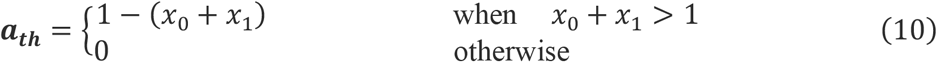

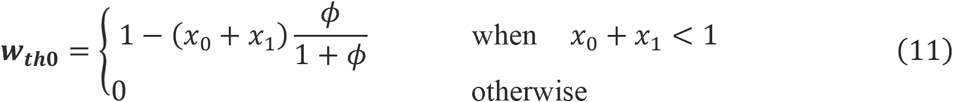

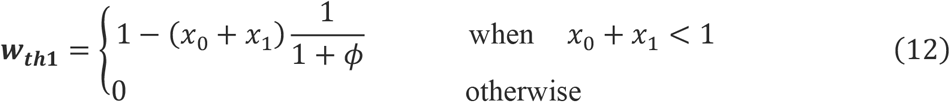

We verified that the key conclusions do not depend on the precise forms of the feedback functions (Supplemental Section C).

Equations 7–12 define the primary model we analyze in this study: a system of two coupled ordinary differential equations with seven parameters. When *x_0_ + x_1_ = 1*, we can further simplify. Taking Equations 7-12 together,

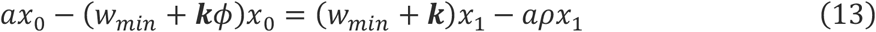

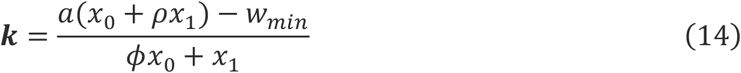

Thus, when *x_0_ + x_1_ = 1*, we have six free parameters: *a, ρ, w_min_, ϕ, m_s_, μ*.

### The model has two regimes. In one regime, neoplastic cells always take over the population

We performed model simulations over a range of parameter values motivated by the experimental literature (detailed in Table S1). We observed two regimes associated with different model behaviors (Figure 2A). Next, we present the corresponding analysis of these two regimes.

**Figure 2.**
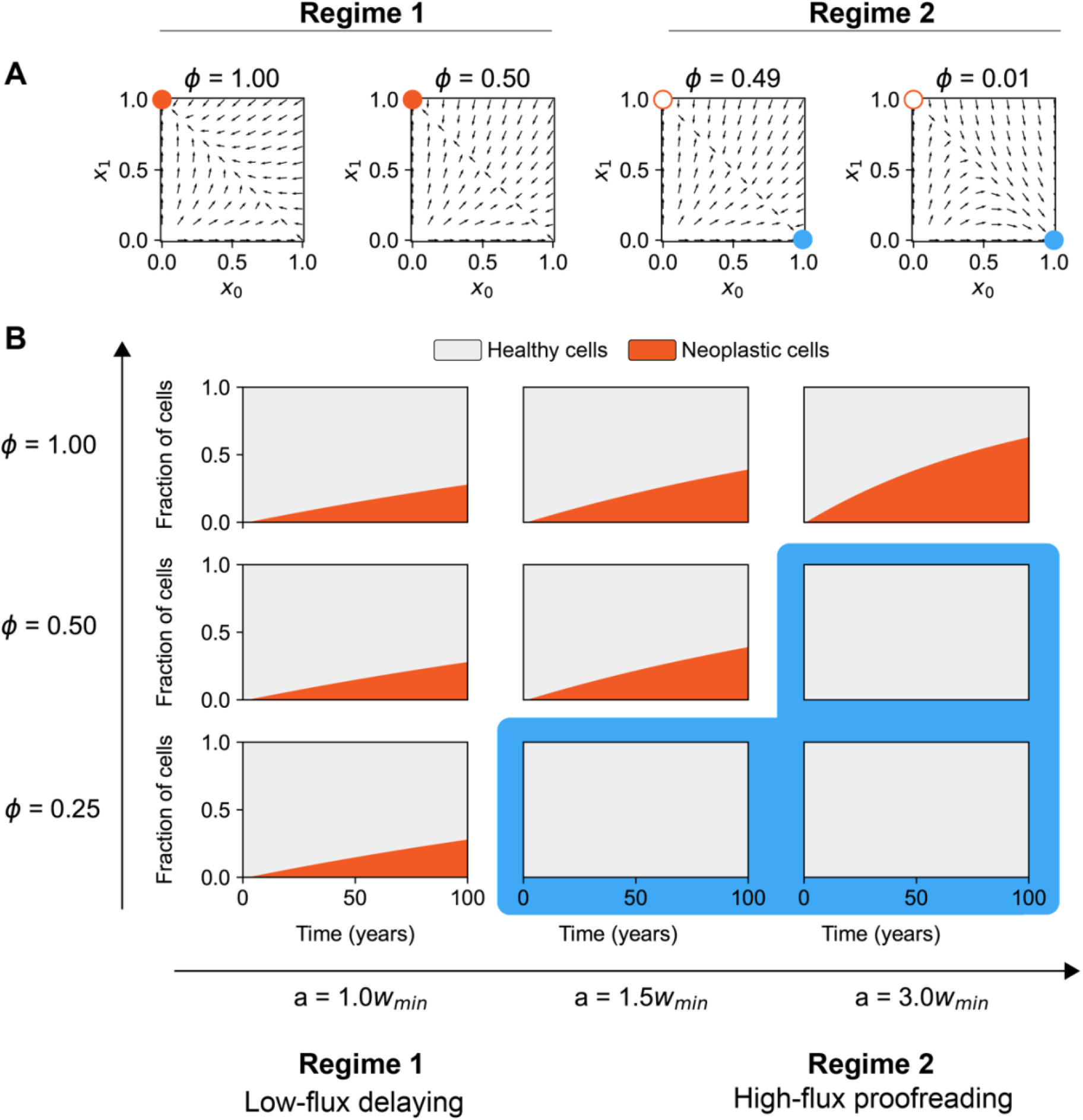
The model solutions fall into two different regimes. **(A)** Phase portraits of the model at different values of *ϕ*. The orange and blue circles indicate fixed points, with filled circles indicating attractors. For these phase portraits, *μ=10^-4^, w_min_ =33.7/year, a =2w_min_, ρ= 1.5, L=1*, and *ϕ* is as indicated. **(B)** Simulations of the model. Orange is the fraction of neoplastic cells. Grey is the fraction of healthy cells. For these simulations, *μ=10^-4^, w_min_ =33.7/year, ρ= 1.5, L=2*, and *a* and *ϕ* are as indicated.

In the first regime, neoplastic cells inevitably take over the population (the orange attractors in Figure 2A; a broader parameter space will be shown in Figure 3). In dynamical systems language, there is only stable fixed point within this regime, corresponding to healthy cells *x_0_ = 0* and neoplastic cells *x_1_* = *1*.

**Figure 3.**
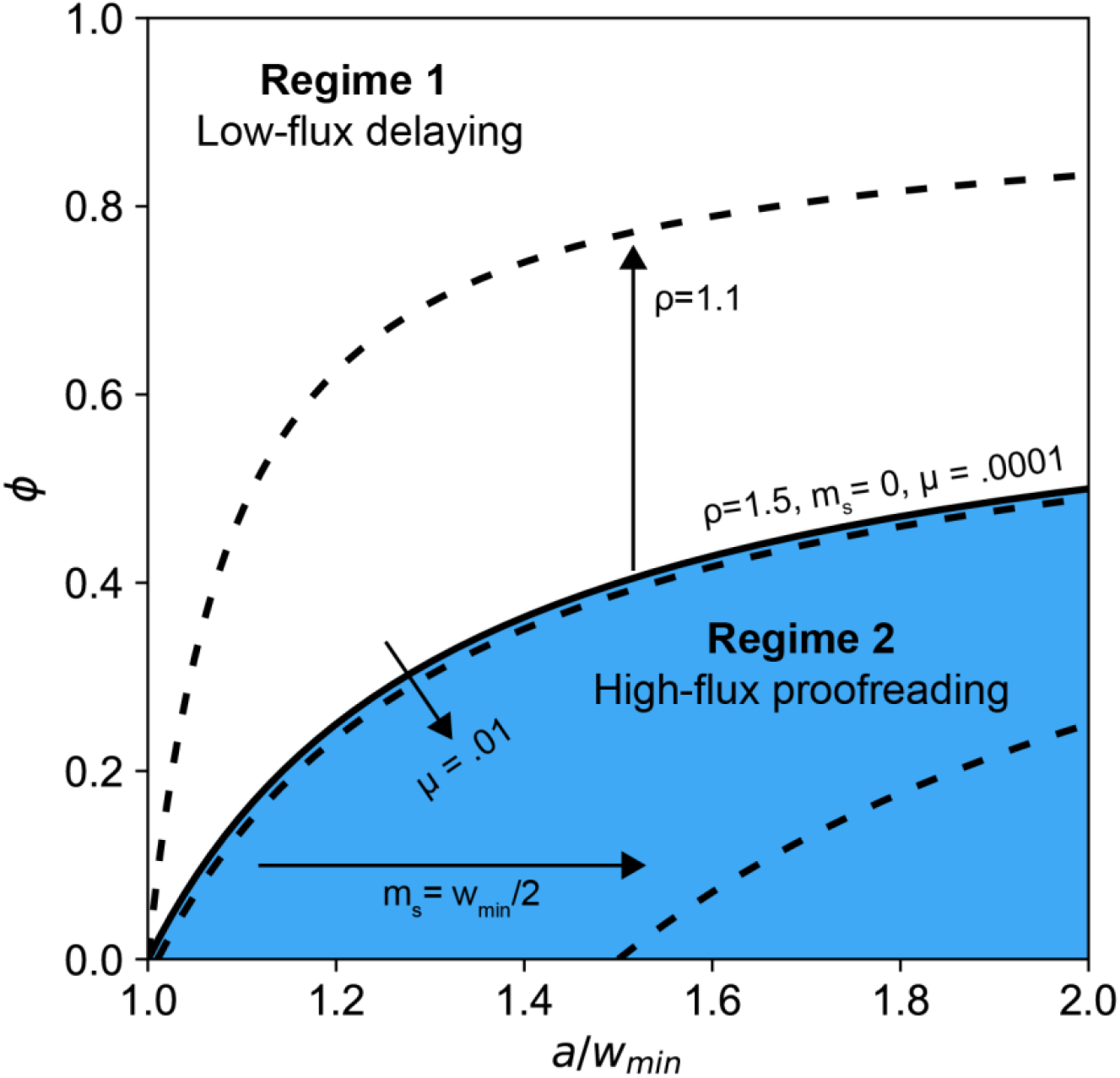
Existence of a regime in the model in which increasing proliferation suppresses accumulation of neoplastic cells. The black solid line indicates the boundary between Regime 1 (white) and Regime 2 (blue) in the model. The analytical expression of the boundary is derived in Condition 17a and plotted here using the parameter values *ρ* = 1.5*, μ* =.0001*, m_s_=*0. Black dashed lines show how the boundary shifts as the parameters vary.

The speed with which neoplastic cells accumulate depends on the parameters, as illustrated in Figure 2B, in which we simulated the model to 100 years (the order of magnitude for human lifespan). In the case where healthy and neoplastic cells divide and die at the same rate (*ρ = 1, ϕ = 1*), we analytically derive the timescale with which neoplastic cells accumulate,

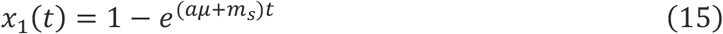

with the initial conditions *x_0_* = 1, *x_1_* = 0. With more than one mutation step *L > 1*,

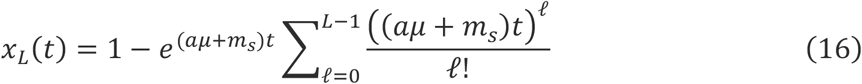

The detailed derivation is presented in Supplemental Sections A and D.

Equations 15-16 show that the speed of accumulation of neoplastic cells *x_1_* increases with proliferation rate *a*, consistent with the intuition that proliferation carries a risk of developing neoplasia. The model behavior in this regime, in which neoplastic cells inevitably take over the population, reproduces previous models of finite populations with genetic drift.^22,23,35,36^ Even without any fitness advantage, a new allele *x_1_* will take over the population because the old allele *x_0_* will drift out. However, the approach to steady state can be slowed by reducing proliferation (Figure 2B, top row). We therefore refer to this strategy as **low-flux delaying**.

### In the second regime of the model, neoplastic cells can be suppressed

Although in one regime of the model the neoplastic cells always overtake the population, in the other regime, neoplastic cells can be suppressed. In simulations, small changes in parameters *a* or *ϕ* moved the attractor from *x_1_=1* to strikingly small numerical values (Figure 2).

To understand when the system can enter this second regime, we derive an exact analytical condition, as detailed in Supplemental Section A:

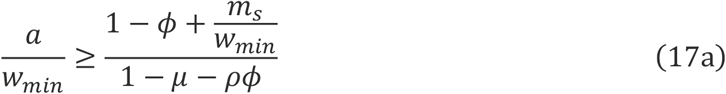

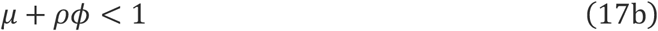

Condition 17a provides a boundary between the two regimes, shown in Figure 3 (solid line). The condition shows that the system can enter the second regime by increasing proliferation *a*. Therefore, surprisingly, increasing proliferation suppresses neoplastic cells. Because proliferation of healthy cells is coupled with proliferation of neoplastic cells (Equation 3), increasing *a* means that the system makes more healthy cells and more neoplastic cells. Because proliferation is coupled with cell death through homeostatic feedback, increasing *a* means that the system also kills more cells, both healthy and neoplastic. As long as there is sufficient discrimination, i.e., the system kills neoplastic cells faster than healthy cells (fast enough to counter their growth advantage *ρ*), the high flux of cells can act as a proofreader to eliminate neoplastic cells. We accordingly call this strategy **high-flux proofreading**.

Increasing flux to eliminate neoplasia may seem costly, but the shape of the boundary between regimes (Figure 3) reveals an intriguing property: increasing flux can be effective even with a seemingly poor detector. Consider *ϕ = 0.5*, a detector that would kill one healthy cell for every two neoplastic cells. The system in Figure 3 can achieve proofreading for *ϕ = 0.5* by increasing *a/w_min_* to slightly more than 2.0, as verifiable with Condition 17. In fact, Condition 17 shows that for *ρ ≈ 1,* the system only needs a slight preference for killing neoplastic cells (*ϕ < 1-μ, μ << 1*) as long as *a/w_min_* is sufficiently high. High-flux proofreading may therefore be a powerful compensatory strategy when a system has limited ability to discriminate between healthy cells and neoplastic cells.

In addition to proliferation and discrimination, the regime in which high-flux proofreading can be achieved also depends on the other parameters in the system (dashed lines in Figure 3). As neoplastic cells gain growth advantage and proliferate faster than healthy cells (*ρ* increases), it becomes harder to suppress their accumulation. Varying the rate at which mutation occurs per cell division (*μ*) has a small effect; increasing the rate of mutation occurring outside cell division (*m_s_*) shifts the boundary to require more proliferation but does not prevent proofreading.

How effective can suppression through high-flux proofreading be? We derive the analytical steady-state solution for *x_1_* in the proofreading regime (Supplemental Section A):

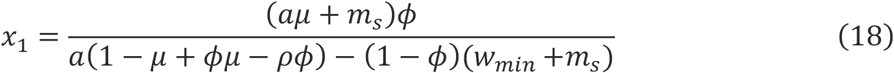

When *L > 1* (Supplemental Section D):

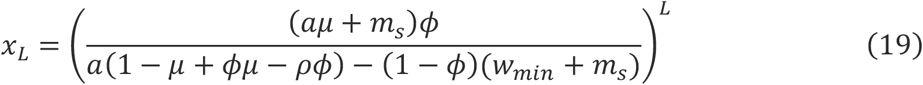

From these equations, we can derive a lower bound on achievable suppression, by taking the limit *a* → ∞ of the solution in Equation 19:

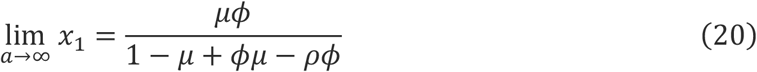

When *ρ = 1*, i.e., when the neoplastic cells have not attained a growth advantage relative to healthy cells, we can use the small-*μ* approximation *μ/(1-μ) ≈ μ*:

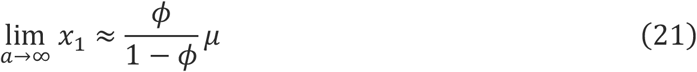

*ϕ/(1-ϕ)* will make *x_1_* smaller for *ϕ < 0.5* and will have only a small effect on order of magnitude for 0.5 < ϕ < 0.9. Therefore, high-flux proofreading can suppress the population of neoplastic cells to a similar order to *μ*. With *L* mutation steps, this can be further decreased to *μ^L^* (Supplemental Section D). To appreciate these parameter values in a quantitative sense, consider that adult humans have on the order of 10^13^ cells.^37^ For *μ* =10^-7^, *L = 2*, as in classical estimates,^25^ high-flux proofreading could in theory suppress the neoplastic population to less than one cell in an organism this size. Further suppression can be achieved with some nonlinear coupling functions (Supplemental Section E), or with models that include stem cells (Supplemental Section F), so these numerical estimates are simply a starting point to highlight that the orders of magnitude described by high-flux proofreading may be in a biologically relevant range.

### Experimental system for assessing high-flux proofreading

The modeling analysis reveals an unexpected strategy in which a high flux of cells acts as a proofreader, eliminating neoplastic cells. Is high-flux proofreading a biologically relevant strategy, as opposed to an esoteric mathematical regime? As a first test, we turned to the moon jelly *Aurelia coerulea*, in which cancer has rarely been observed.^20,21^ We reason that a cancer-resistant animal could provide a clear first test of the model prediction.

In *Aurelia*, we focused on the juvenile stage (called ephyra; Figure 4A) because at this stage the animal displays a striking and stereotypic eight-fold symmetry of the discrete appendages, making assessment of abnormal growths (neoplasms) straightforward. We did not assess for the invasiveness of neoplasms because distinguishing neoplasia and cancer is outside the scope of the current model predictions.

**Figure 4.**
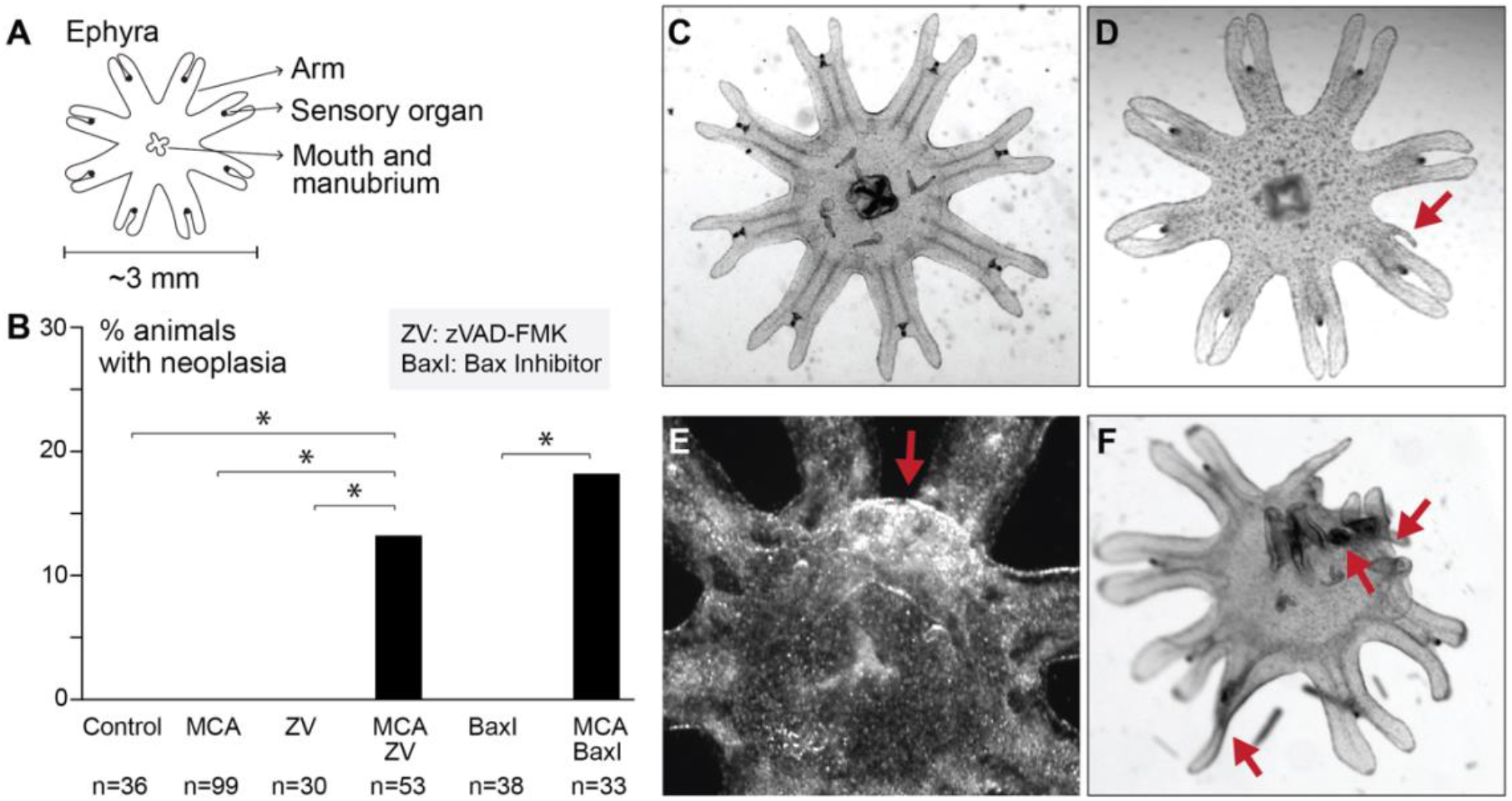
*Aurelia* ephyrae can be induced to produce neoplasms. **(A)** We performed the experiments in the juvenile moon jelly, called the ephyra. The ephyra has a radially symmetrical body, with eight symmetrically distributed appendages. **(B)** In these experiments, ephyrae were exposed to one or a combination of the following: no treatment (Control), MCA (330 μM and 500 μM), zVAD-FMK (ZV; 500 μM), Bax-inhibiting peptide (BaxI; 100 μM). The ephyrae were monitored for up to 7 days. Each dataset came from 2–9 independent experiments, with the total number of ephyrae examined shown below the relevant column (n). Statistical significance was assessed using Fisher’s exact test (p <0.05). **(C)** A typical ephyra in artificial salt water, showing no abnormal morphology. **(D-F)** Co-treatment of carcinogen with apoptosis inhibitor produced neoplasms (red arrows). Figure 4E was taken in darkfield to better capture the abnormal growth.

As a carcinogen, we used 3-methylcholanthrene (MCA), a known carcinogen in humans and rodents.^38–40^ Treating the ephyrae with MCA, at ∼20 times the doses that induce cancer formation in mice (see Methods), did not produce any morphological abnormalities for experiments up to seven days in duration (Figure 4B). Thus, as expected from the rare observation of cancers in cnidaria, *Aurelia* ephyrae resist a high dose of carcinogen.

Next, having established that *Aurelia* is resistant to carcinogens, we assessed expected ways with which experiments can break this resistance. A considerable experimental literature supports the idea that apoptosis mediates the targeted killing of neoplastic cells.^8,41,42^ To inhibit apoptosis, we treated the ephyrae with a caspase inhibitor (zVAD-FMK), which has previously been shown to be effective in *Aurelia* ephyrae.^41^ Co-treating with carcinogen and zVAD-FMK produced neoplasms in 13% of ephyrae (Figure 4B). Neoplasms could be detected as rapidly as within a day of treatment. The degree of neoplasia varied, from a small growth on the appendage (Figure 4D), to a large lump in the body (Figure 4E), to in the most dramatic cases, a gross disruption of symmetry and morphology (Figure 4F). To verify that the effect was due to blocking apoptosis, as opposed to non-specific effects of the chemical used, we tested another apoptosis inhibitor, a Bax-inhibiting peptide. As with zVAD-FMK, co-treatment with MCA and Bax inhibition produced neoplasms in 18% of ephyrae (Figure 4B). Neither zVAD-FMK nor Bax inhibition alone produced neoplasms (Figure 4B). These experiments demonstrate, as expected, that activation of apoptosis mediates suppression of neoplasia in *Aurelia*.

### *Aurelia* increases cell proliferation in response to carcinogen treatment

Because neoplasms could be induced in *Aurelia*, we used this controlled experimental system to test a novel hypothesis. The mathematical model predicts that, in response to carcinogen, a high-flux proofreading system will increase proliferation as part of its protective response. To test this prediction, we assessed cell proliferation using 5-ethynyl-2′-deoxyuridine (EdU), a thymidine analog that is incorporated into newly synthesized DNA.^43^ Carcinogen-treated ephyrae show strikingly elevated EdU signals (Figure 5). This assay demonstrates that *Aurelia* increases cell proliferation in response to carcinogen treatment.

**Figure 5.**
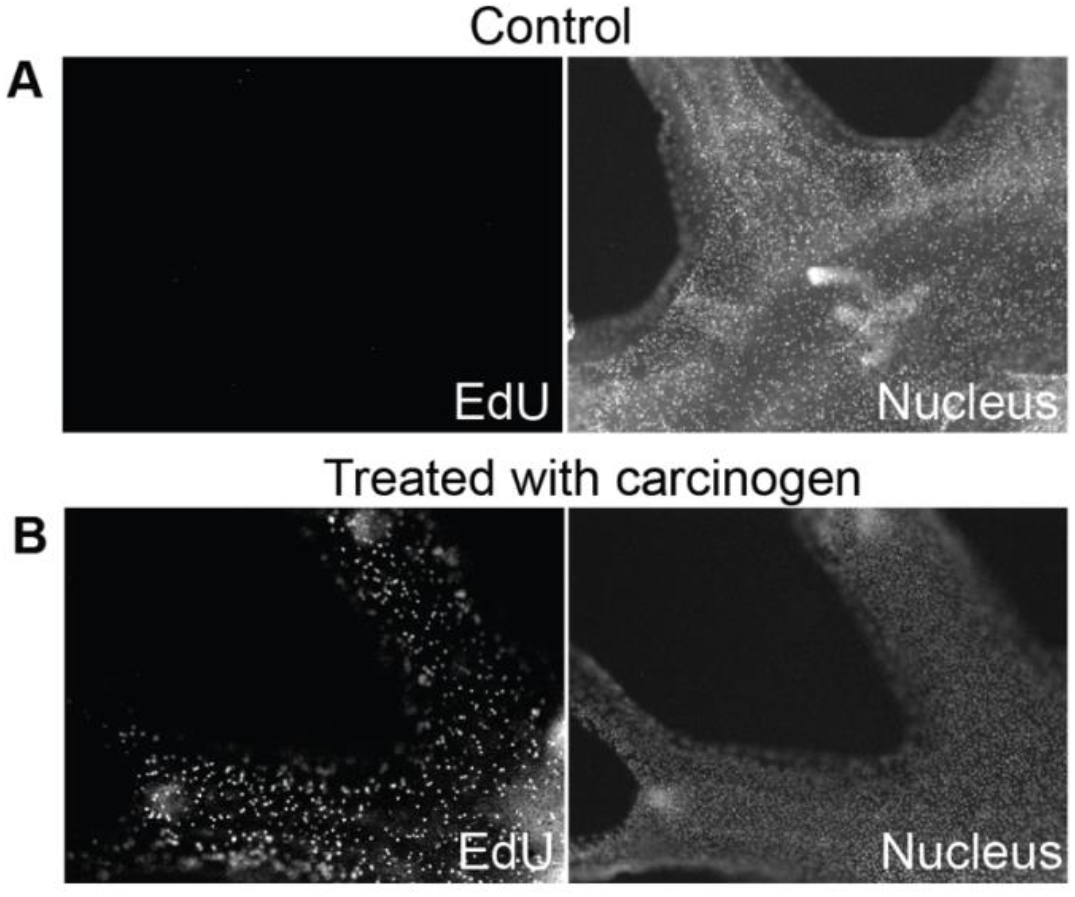
Cell proliferation increases upon carcinogen exposure. Control or carcinogen-treated ephyrae were fed with EdU for 8 hours and then fixed for staining. As a positive control, the ephyrae were stained with Hoechst to mark the nuclei. **(A)** Control ephyrae show negative EdU signals (n=4). **(B)** Ephyrae treated with 500 μM carcinogen MCA show markedly elevated EdU signals (n=4).

### Inhibiting proliferation potentiates neoplasia, while rescuing proliferation rescues resistance

The increased proliferation upon carcinogen exposure is consistent with the model prediction that the organism increases proliferation as part of the cancer-protective response. However, the observation can also be consistent with an alternative explanation that the carcinogen itself increases proliferation when it produces neoplasms. To assess whether increased proliferation protects against neoplasia, we tested inhibiting proliferation. First, we leveraged physiological cues that are known to inhibit proliferation in *Aurelia*, specifically stress.^44^ Ephyrae exposed to osmotic stress and then treated with carcinogen (MCA) showed markedly decreased proliferation compared to those treated with MCA alone (Figure 6B). Consistent with the hypothesis that the increased proliferation is protective, neoplasms were observed in 15% of exposed to osmotic stress and treated with carcinogen (Figure 6A). As another way to produce stress, we tested injury.^44,45^ When injured ephyrae were treated with carcinogen, 20% of the ephyrae developed neoplasms (Figure 6A). As a negative control, we verified that stressed ephyrae, with osmotic stress or injury alone, showed no neoplasms (Figure 6A).

**Figure 6.**
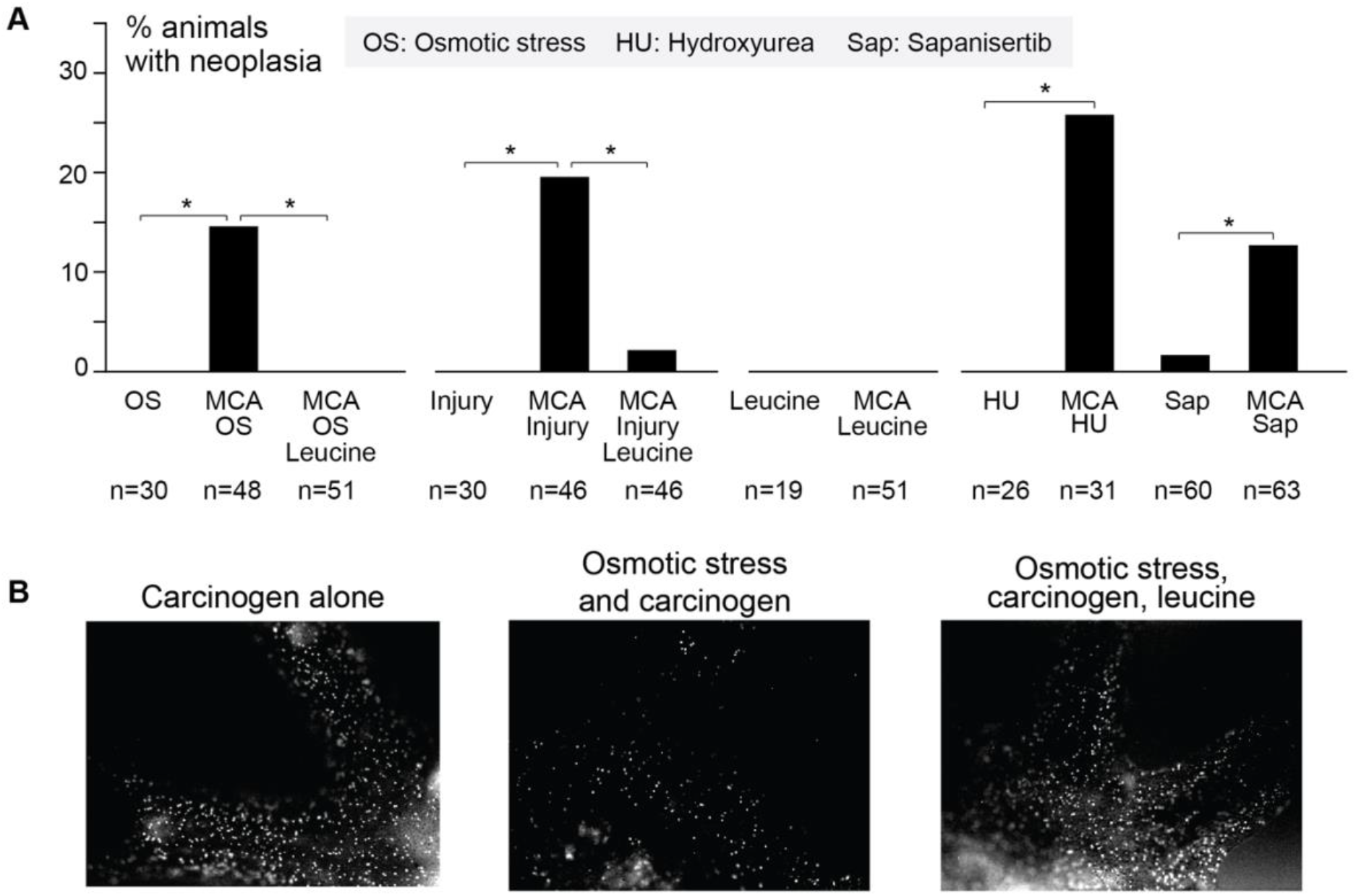
Modulating proliferation modulates resistance to neoplasia. **(A)** In these experiments, osmotic stress was implemented by diluting the concentration of artificial salt water by half for thirty minutes. Injury was implemented by amputation of three appendages. For the treatment, ephyrae were placed in one or a combination of the following: 500 μM MCA,10 μM hydroxyurea (HU), 1 nM sapanisertib (Sap), or 100 μM leucine. Ephyrae were examined for abnormal growths for 1–7 days after the start of treatment. Statistical significance was assessed using the Fisher exact test (p<0.05). **(B)** Ephyrae were fed with EdU, which marks proliferating cells. Left to right: An ephyra treated with 500 μM MCA (n=4), reproduced from Figure 5B; an ephyra exposed to osmotic stress and then treated with 500 μM MCA (n= 3); an ephyra exposed to osmotic stress and then treated with 500 μM MCA and 100μM leucine (n= 3).

Stress or injury is a broad stimulus that can plausibly potentiate neoplasia in other ways than inhibiting proliferation. To verify the protective effects of inhibiting proliferation, we tested known small-molecule inhibitors of proliferation. Hydroxyurea is an antiproliferative drug that has been used in the treatment of some cancers.^46^ Hydroxyurea treatment alone did not lead to neoplasms (Figure 6A). Co-treating with carcinogen and hydroxyurea led to neoplasms in 26% of ephyrae (Figure 6A). To further verify that inhibiting proliferation potentiates neoplasms, we tested sapanisertib, another antiproliferative agent.^47^ Sapanisertib alone led to a neoplasm in one ephyra out of 60 (2%; Figure 6A). Co-treatment with MCA and sapanisertib led to significantly more ephyrae with neoplasms (13%; Figure 6A). The low but nonzero rate of neoplasms with sapanisertib alone is consistent with the idea that proofreading is a physiological function in the setting of routine, spontaneous mutation, even without carcinogenic treatment. Together, these data demonstrate that blocking proliferation, with physiological cues or antiproliferative drugs, reduces resistance to neoplasia.

To further verify that proliferation protects against neoplasia, we performed rescue experiments. We reasoned that if proliferation is indeed protective, then restoring proliferation should rescue resistance to neoplasia. It has been shown that increasing feeding can rescue growth in stressed ephyrae, and that this effect can recapitulated by the branched-chain amino acid leucine.^45^ Feeding leucine to the stressed ephyrae indeed, as expected, promoted proliferation (Figure 6B). Consistent with the hypothesis that proliferation is protective, feeding leucine to osmotically stressed ephyrae rescued resistance to neoplasia (to 0%; Figure 6A). Similarly, feeding leucine to injured ephyrae rescued resistance (to 2%; Figure 6A). Thus, inhibiting proliferation in *Aurelia*, through drugs or physiological stress, potentiates neoplasia, and restoring proliferation through leucine feeding rescues resistance to neoplasia.

## Discussion

In this study, we developed a general model of cell birth, death, and mutation. We used the model to assess conditions that suppress accumulation of neoplastic cells while also maintaining tissue homeostasis. We find that, although increasing proliferation can increase the risk of accumulating neoplastic cells, there is a regime in the model in which increasing proliferation instead suppresses accumulation of neoplastic cells. In this regime, the system makes more and kills more cells as a way to proofread neoplastic cells. We call this high-flux proofreading. As a first experimental test, we investigated high-flux proofreading in the moon jelly. Neoplasms have rarely been observed in cnidarians, and yet simply inhibiting proliferation is sufficient to promote neoplasms in the moon jelly. Together, the modeling and experimental analyses here identify high-flux proofreading as an effective cancer resistance strategy.

The processes in our model are conserved cellular dynamics that are not specific to moon jellies. Accordingly, the model could be applied across animal systems. For instance, two well-known examples of mammals with low rates of cancer are naked mole rats and elephants. These systems use increased sensitivity to mutation or crowding to trigger apoptosis and target potential cancers.^8,10,14,42,48–50^ In the model, it is possible to make the system more sensitive by increasing proliferation *a* for a given detector (fixed *ϕ*), or by improving detection (decreasing *ϕ*). The coupling functions themselves (e.g., linear or nonlinear) could also vary across animals. Thus, the model offers a framework that could be useful for quantitatively defining how cancer suppression is achieved, and for comparing how evolution may tune parameters across animal species.

High-flux proofreading may already be operating invisibly, and effectively, as part of tissue homeostasis in humans. Because capacity for tissue maintenance reduces as we age, for instance with accumulation of senescent cells or fibrous tissue, high-flux proofreading may become less effective with age. High-flux proofreading may also fail due to mutations that take the system out of the regime in which it is effective. In the model, high-flux proofreading can fail with sufficient loss of DNA repair (increase in *μ*), sufficient growth advantage for neoplasia (increase in *ρ*), and sufficient evasion of targeted apoptosis (increase in *ϕ*). These are indeed conditions that are observed in clinically relevant cancers.^41^ Therefore, the modeling framework can serve as the basis for a more translational model to inform, e.g., early treatments of precancerous cells, prevention of secondary carcinogenesis as a complication of anti-proliferative treatments, or high flux as an augmentative strategy for immunotherapies. High flux may also be a useful guiding principle for regenerative therapies that use stem cells, for which unrestrained and neoplastic growth is a potential risk.^51–53^

There is an old engineering rule of thumb that a technology can achieve only two of the three goals of quality, speed, and low cost. In high-flux proofreading, the system achieves quality control and responsiveness to demand (speed) at the cost of a high flux of cells. This approach to quality control appears to be a recurring strategy in biology. For example, in cell signaling, it is common for the signal transducer to be continuously produced and degraded, a seemingly futile cycle that non-intuitively gives rise to robustness in signaling.^54,55^ In protein regulation, the number of proteins can theoretically be regulated, despite the inherent stochasticity of transcription, through overproduction and overdegradation.^56^ In the immune system, immature T cells are considerably overproduced, with over 95% of cells dying in thymic selection to achieve quality control in the T cell receptor repertoire.^57,58^ And lastly, for biosynthetic reactions, Hopfield showed that kinetic proofreading (the use of high-energy intermediates with seemingly wasteful side reactions) improved the accuracy of protein transcription or DNA replication.^59^ There may be similar underlying mathematical structures between Hopfield’s kinetic proofreading and the high-flux proofreading proposed in this study; in both models, increasing the number of intermediate steps geometrically decreases the error rate. Thus, from molecules to cells to tissues, a richer theory of this type of control in biology may uncover deeper constraints that shape living systems.

## Acknowledgements

The authors thank Michael Abrams, Ty Basinger, and Aki Ohdera for guidance in jellyfish care and imaging, and Yutian Li for help with imaging. This work was supported by the Army Research Office Multidisciplinary University Research Initiative (W911NF-17-1-0402 to JCD) and the Carver Mead New Adventures Fund (JCD and LG).

## Author Contributions

AAS: Conceptualization, Data curation, Formal analysis, Investigation, Methodology, Software, Validation, Visualization, Writing – original draft, Writing – review & editing

LG: Conceptualization, Funding acquisition, Investigation, Methodology, Project administration, Resources, Supervision, Validation, Visualization, Writing – original draft, Writing – review & editing

JCD: Conceptualization, Funding acquisition, Investigation, Methodology, Project administration, Supervision, Writing – review & editing

## Material Availability

Code used to make figures is available at https://github.com/anishsarma/proofreading.

## Materials and Methods

### Mathematical modeling

Numerical results and figures were produced in Python, using a Jupyter notebook with packages Pandas, NumPy, SciPy, and MatPlotLib.

### Jellyfish care and strobilation

*Aurelia coerulea* polyps and ephyrae were maintained in 32 ppt artificial salt water (Instant Ocean mix) at 68°F on a 12-hour light cycle. Strobilation was induced by incubation in 50 mM 5-methoxy-2-methyl-indole (VWR 101844-752) for an hour. Strobilated ephyrae were fed with rotifers (*Brachionus plicatilis*, Reed Mariculture) or brine shrimp (*Artemia nauplii*, Brine Shrimp Direct) every 2–3 days.

### Chemical treatment

Ephyrae were screened for normal morphology (symmetric, 8-10 lobes, no inter-lobe nodules, no bell formation, planar) prior to inclusion in the experiments. Ephyrae were placed in a 12-or 24-well plate, at 8–12 ephyrae per well and then incubated in molecules or chemicals (in a total 1-2 mL). Plates were rocked at a rate of ∼0.5 Hz. Ephyrae were not fed during treatment. The following chemicals were used: 3-methylcholanthrene (VWR 89158-830, 89158-832); z-VAD-FMK (Fisher Scientific PRG7231, PRG7232, 50202909; VWR 103521, 89146-938); Sapanisertib (Fisher Scientific 502028984, 501362995; VWR 103542-016); Hydroxyurea (Sigma-Aldrich B2261; Life Technologies 151680050); Bax-Inhibiting Peptide (Fisher Scientific 1968115MG, 80603-646); L-leucine (VWR E811). Abnormal growths, as described in Figure 4D-F, were assessed at 1–7 days after the start of treatment. The MCA dose we used in the jellyfish is ∼20 times of that used in the mice, estimated in this way: MCA dose used in mouse studies ranges from 0.005–0.05 mg per gram of animal weight. In this study, each jelly was exposed to ∼15 ug of MCA (at 500 μM concentration). Using a typical dry weight of *Aurelia* of ∼100–1000 μg,^60^ and assuming that soft marine animals readily absorb environmental small molecules, the dose of MCA used in this study is ∼0.1–1 mg per gram of animal weight.

### Injury and Osmotic Stress

Amputation injury was performed following methods described previously.^44,45^ Ephyrae were cut across the body removing three lobes and part of the bell using a razor blade. Osmotic stress was induced by incubating the ephyrae in ASW diluted to 50% (16 parts per thousand) by deionized water, for 30 minutes. After the osmotic stress, ephyrae were returned to typical ASW and then immediately transferred to the experimental well for chemical treatment.

### Fluorescent staining

To image proliferating cells, ClickIT EdU AlexaFluor488 (Fisher Scientific, C10337) was used, following a protocol that has previously been established in ephyrae.^44^ For these experiments, osmotically stressed ephyrae were pre-stressed as described above. Ephyrae were incubated in 1:1000 EdU and artificial salt water along with the relevant chemical treatment for an 8-hour treatment period. Following the existing protocol,^44^ ephyrae were then fixed (3.7% formaldehyde/PBS), permeabilized (0.5% Triton X-100/PBS), and incubated in the dark with the ClickIT reaction mixture. Lastly, nuclei were stained with Hoechst 33342 (Sigma-Aldrich B2261).

### Imaging

Brightfield, darkfield, and fluorescent images of jellyfish were acquired using a Zeiss AxioZoom.V16 stereo zoom microscope.

