## Supplementary Materials for "Proliferation to suppress neoplasia: a general model and a first test in the moon jelly"

**A. Deriving analytical solutions of the model**

The basic model with $L = 1$, as described in the main text, is:

$$\frac{dx_0}{dt} = \boldsymbol{a_0}x_0 - \boldsymbol{w_0}x_0 - \boldsymbol{m}x_0 \tag{A1}$$

$$\frac{dx_1}{dt} = \boldsymbol{a_1}x_1 - \boldsymbol{w_0}x_1 + \boldsymbol{m}x_0 \tag{A2}$$

**Tissue homeostasis.** Tissue homeostasis is achieved when the total population size is $x_0 + x_1 = 1$ and stable:

$$\frac{dx_0}{dt} + \frac{dx_1}{dt} = 0 \quad \text{when} \quad x_0 + x_1 = 1 \tag{A3a}$$

$$\frac{dx_0}{dt} + \frac{dx_1}{dt} < 0 \quad \text{when} \quad x_0 + x_1 > 1 \tag{A3b}$$

$$\frac{dx_0}{dt} + \frac{dx_1}{dt} > 0 \quad \text{when} \quad x_0 + x_1 < 1 \tag{A3c}$$

Summing Equations A1 and A2, we find the summed derivative:

$$\frac{dx_0}{dt} + \frac{dx_1}{dt} = \boldsymbol{a_0}x_0 - \boldsymbol{w_0}x_0 - \boldsymbol{m}x_0 + \boldsymbol{a_1}x_1 - \boldsymbol{w_1}x_1 + \boldsymbol{m}x_0 \tag{A4}$$

Therefore, the conditions that are required for tissue homeostasis are:

$$\boldsymbol{a_0}x_0 - \boldsymbol{w_0}x_0 + \boldsymbol{a_1}x_1 - \boldsymbol{w_1}x_1 = 0 \quad \text{when} \quad x_0 + x_1 = 1 \tag{A5a}$$

$$\boldsymbol{a_0}x_0 - \boldsymbol{w_0}x_0 + \boldsymbol{a_1}x_1 - \boldsymbol{w_1}x_1 < 0 \quad \text{when} \quad x_0 + x_1 > 1 \tag{A5b}$$

$$\boldsymbol{a_0}x_0 - \boldsymbol{w_0}x_0 + \boldsymbol{a_1}x_1 - \boldsymbol{w_1}x_1 > 0 \quad \text{when} \quad x_0 + x_1 < 1 \tag{A5c}$$

Solving Equation A5a gives us a tissue homeostasis constraint that must hold for any rate functions as long as $x_0 + x_1 = 1$:

$$\boldsymbol{a_0}x_0 - \boldsymbol{w_0}x_0 = \boldsymbol{w_1}x_1 - \boldsymbol{a_1}x_1 \tag{A6}$$

We want to understand how the relationships between the rate functions ($\boldsymbol{a_0}$, $\boldsymbol{w_0}$, $\boldsymbol{a_1}$, and $\boldsymbol{w_1}$) shape the model behavior. In this section, in order to derive analytical solutions, we assume linear coupling relationships. Later, we generalize the findings to the cases of nonlinear couplings (Section E).

**Linear coupling relationships.** We will use the following relationships between the rate functions:

Mutation function:

$$\boldsymbol{m} = \boldsymbol{a_0}\mu + m_s \tag{A7}$$

Proliferation function:

$$\boldsymbol{a_1} = \rho \boldsymbol{a_0} \quad \text{(A8)}$$

Death functions:

$$\boldsymbol{w_0} = w_{min} + \boldsymbol{k}\phi \quad \text{(A9a)}$$
$$\boldsymbol{w_1} = w_{min} + \boldsymbol{k} \quad \text{(A9b)}$$

**Regime 1: Low-flux delaying**

We start with the simple case in which neoplastic cells and healthy cells divide and die at the same rate, i.e., $\rho = 1$, $\phi = 1$, so that $\boldsymbol{w_0} = \boldsymbol{w_1} = \boldsymbol{w}$ and $\boldsymbol{a_0} = \boldsymbol{a_1} = \boldsymbol{a}$. The tissue homeostasis constraint in Equation A6 simplifies to:

$$\boldsymbol{a}x_0 - \boldsymbol{w}x_0 = \boldsymbol{w}x_1 - \boldsymbol{a}x_1 \quad \text{(A10)}$$
$$\boldsymbol{a}(x_0 + x_1) = \boldsymbol{w}(x_0 + x_1) \quad \text{(A11)}$$

And because $x_0 + x_1 = 1$ when the tissue homeostasis constraint is satisfied,

$$\boldsymbol{a} = \boldsymbol{w} \quad \text{(A12)}$$

Therefore, the tissue homeostasis constraint can be satisfied if $\boldsymbol{a}$ and $\boldsymbol{w}$ are constants along the line $x_0 + x_1 = 1$, with feedback functions off of the line as described in main text Equations 10-12. We now make this further simplification:

$$a = w \quad \text{(A13)}$$

Substituting Equation A13 and Equation 7 back into Equation A2, we can express the dynamics along the line $x_0 + x_1 = 1$ entirely in terms of $x_1$:

$$\frac{dx_1}{dt} = ax_1 - wx_1 + (a\mu + m_s)x_0 \quad \text{(A14)}$$

Equation A15 has steady states where $a\mu + m_s = 0$ or where $x_1 = 1$. $a\mu + m_s = 0$ requires $a = 0$ and $m_s = 0$. Biologically, these conditions would mean that the tissue has zero turnover ($a$=$w$=0) and faces no exposure-driven mutations ($m_s$). Outside of this special case, the steady state is $x_1 = 1$: the neoplastic cells will always eventually overtake the population.

From Equation A15, we find a closed-form solution for $x_1(t)$ from the initial conditions $x_0 = 1$, $x_1 = 10$

$$x_1(t) = 1 - e^{(a\mu + m_s)t} \quad \text{(A16)}$$

Although the steady state is always $x_1 = 1$, the rate at which this steady state is approached can be reduced by reducing proliferation (making $a$ small).

**Regime 2: High-flux proofreading**

We return to the basic model (Equation A1-2), using the linear mutation function as defined in Equation A7:

$$\frac{dx_0}{dt} = \boldsymbol{a_0}x_0 - \boldsymbol{w_0}x_0 - (\boldsymbol{a_0\mu} + m_s)x_0 \tag{A17}$$

$$\frac{dx_1}{dt} = \boldsymbol{a_0}x_1 - \boldsymbol{w_0}x_1 + (\boldsymbol{a_0}\mu + m_s)x_0 \tag{A18}$$

In the previous section, we considered the case that in which neoplastic cells and healthy cells divide and die at the same rate ($\rho = 1$ and $\phi = 1$), and found that in this condition, there is only one steady state possible, i.e., $x_1 = 1$. We now consider the case $\phi < 1$: neoplastic cells die at a higher rate than healthy cells because of detection-driven apoptosis. Intuitively, if neoplastic cells die fast enough, an additional steady state could exist with $x_1 << 1$.

To look for the steady states in the system, we set Equations A17–A18 to zero. At steady state, since $x$ is constant, the rate functions also become constants.

$$0 = a_0x_0 - w_0x_0 - (a_0\mu + m_s)x_0 \tag{A19}$$

$$0 = a_0x_1 - w_0x_1 + (a_0\mu + m_s)x_0 \tag{A20}$$

There are three possibilities associated with Equations A19-A20.

1. $x_0 = 0$. Equation A19 can be satisfied if $x_0$ is zero (meaning $x_1 = 1$). In that case, to satisfy Equation A20, we require $a_1 = w_1$. Note that although the tissue homeostasis in Equation A6 constraint allows $\boldsymbol{a_1}$ and $\boldsymbol{w_1}$ to vary with $x$, it requires $\boldsymbol{a_1} = \boldsymbol{w_1}$ when $x_1 = 1$. (To see this, substitute $x_0 = 0$ and $x_1 = 1$ into Equation A6.) Therefore, as long as tissue homeostasis is achieved, a steady state always exists at $x_1 = 1$.
2. $x_0 < 0$ or $x_1 < 0$. No steady state exists that satisfies the existing constraints. This includes not only the explicit tissue homeostasis constraint (Equation A6) and the associated constraint $x_0 + x_1 = 1$ but also the nonnegativity constraints implicit to any model of quantities like populations: $x_0, x_1$, and all rate functions must be nonnegative. For example, if $a_1 > w_1$, Equation A20 can only be satisfied by $x_0 < 0$. The biological intuition for this example is that if $a_1 > w_1$, $x_1$ should grow exponentially, and tissue homeostasis will not be achieved.
3. $x_0 > 0$. If $x_0$ is nonzero and positive (meaning $x_1 < 1$), then we can divide both sides of Equation A19 by $x_0$ to solve for an additional steady state. If we can derive a valid $x_1 < 1$ (nonnegative, with all rates nonnegative and $x_0$ nonnegative), then the steady state exists in addition to the already established steady state at $x_1 = 1$. If we cannot derive a valid $x_1 < 1$, but tissue homeostasis is still achieved, then the steady state at $x_1 = 1$ still exists.

Pursuing the third possibility with nonzero $x_0$, we divide both sides of Equation A19 by $x_0$.

$$0 = a_0 - w_0 - (a_0\mu + m_s) \tag{A21}$$
$$a_0 - w_0 = a_0\mu + m_s \tag{A22}$$

Now, substituting Equation A21 into Equation A17,

$$0 = a_1x_1 - w_1x_1 + (a_0 - w_0)(1 - x_1) \tag{A23}$$

Solving for $x_1$, we obtain the analytical expression describing the steady-state neoplastic population.

$$x_1 = \frac{a_0 - w_0}{w_1 - a_1 - (w_0 - a_0)} \tag{A24}$$

To build intuition for the conditions that enable neoplasia suppression via upregulating apoptosis, consider the case when the healthy and neoplastic cells proliferate at the same rate (using Equation A8 with ρ = 1, set $a_0 = a_1 = a$):

$$x_1 = \frac{a - w_0}{w_1 - w_0} \tag{A25}$$
$$x_1 = \frac{\frac{a}{w_0} - 1}{\frac{w_1}{w_0} - 1} \tag{A25}$$

We see that the more discrimination the system can achieve, i.e., the larger $w_1/w_0$, the smaller the steady-state $x_1$ becomes. In the special case where $\phi = 0$,

$$x_1 = \frac{a - w_{min}}{k} \tag{A25}$$

Next, to explicitly derive the dependence of $x_1$ on $a$, we start with the steady-state solution found in Equation A24. We substitute proliferation coupling from Equation A8:

$$x_1 = \frac{a - w_0}{w_1 - \rho a - (w_0 - a)} \tag{A28}$$

From Equation A21, we know that $w_0 = a(1 - \mu) + m_s$. Substituting this expression into Equation A24, we find:

$$x_1 = \frac{a - a(1 - \mu) + m_s}{w_1 - \rho a - (a - a(1 - \mu) + m_s)} \tag{A29}$$

Next, we find $w_1$ in terms of $w_0$. If $\phi = 0$, then we can always select a $w_1$ that makes $x_1$ arbitrarily small. If $\phi > 0$, then from Equation A9a:

$$k = \frac{w_0 - w_{min}}{\phi} \tag{A30}$$

Substituting this into Equation A9b,

$$w_1 = w_{min} + \frac{w_0 - w_{min}}{\phi} \tag{A31}$$

We then rearrange Equation A21 and substitute:

$$w_1 = w_{min} + \frac{a(1-\mu) - m_s - w_{min}}{\phi} \tag{A32}$$

$$w_1 = \frac{a(1-\mu) - m_s - (1-\phi)w_{min}}{\phi} \tag{A33}$$

Substituting Equation A33 into Equation A29, we solve for $x_1$ in terms of $a$:

$$x_1 = \frac{a - a(1-\mu) + m_s}{\frac{a(1-\mu) - m_s - (1-\phi)w_{min}}{\phi} - \rho a - (a - a(1-\mu) + m_s)} \tag{A34}$$

Simplifying:

$$x_1 = \frac{(a\mu + m_s)\phi}{a(1 - \mu + \phi\mu - \rho\phi) - (1-\phi)(w_{min} + m_s)} \tag{A35}$$

In Equation A35, proliferation $a$ appears both in the numerator and the denominator. Depending on the relative magnitudes of the other parameters, there is a regime in the model in which increasing proliferation $a$ suppresses the steady-state population of neoplastic cells $x_1$.

Taking the limit:

$$\lim_{a\to\infty} x_1 = \frac{\mu\phi}{1 - \mu + \phi\mu - \rho\phi} \tag{A36}$$

Equation A36 shows that the steady-state neoplastic cell population $x_1$ *increases* with the false positive rate of the apoptotic detector $\phi$; *increases* with the per-division mutation rate $\mu$; and *increases* with the neoplastic growth advantage $\rho$. Most importantly, the steady-state neoplastic cell population *decreases* with proliferation $a$.

Two interesting special cases are $\rho = 0$ and $\rho = 1$. $\rho = 0$ describes the case where control of neoplasia is mediated through perfect discrimination in division. Even as $\rho$ approaches zero, $x_1$

approaches a nonzero value. By contrast, when $\phi$ approaches zero, $x_1$ approaches zero. This suggests that discrimination through cell death is more effective than discrimination through cell division. $\rho = 1$ describes the case where neoplastic cells, early in transformation, have not attained any growth advantage relative to healthy cells. This case allows some additional simplification to highlight the key parameters that govern the system behavior early in neoplastic transformation. In particular, for small per-division mutation rate $\mu$, we use the Taylor approximation $\mu/(1-\mu) \approx \mu$, so that Equation A36 is well-approximated by:

$$\lim_{a\to\infty} x_1 \approx \frac{\mu\phi}{1-\phi} \tag{A37}$$

These equations offer intuition into how $x_1$ varies with key functions, especially proliferation, mutation, and discrimination. However, Equations A36-A37 are specific to linear coupling functions. With nonlinear coupling functions, lower values of $x_1$ can be obtained in the high-proliferation limit, to a lower bound of zero.

**Boundary between the regimes (Figure 3)**

We next want to establish the exact conditions under which suppression is achieved. Taking the right-hand side of Equation A36:

$$0 \leq \frac{\mu\phi}{1-\mu+\phi\mu-\rho\phi} \leq 1 \tag{A38}$$

To multiply through by the denominator without changing the direction of the inequalities, the denominator must be positive (as it must be for any nonnegative steady state, because each constant is nonnegative and the numerator is nonnegative). Therefore, we split Condition A38 into two conditions:

$$0 \leq \mu\phi \leq 1-\mu+\phi\mu-\rho\phi \tag{A39a}$$
$$0 \leq 1-\mu+\phi\mu-\rho\phi \tag{A39b}$$

If Condition A39a is met, Condition A39b is necessarily met, so we consider Condition A39a alone and simplify.

$$0 \leq 1-\mu-\rho\phi \tag{A40}$$

Stating this in terms of how effective discrimination needs to be, with smaller $\phi$ associated with better discrimination in the limit with high $a$:

$$\phi \leq \frac{1-\mu}{\rho} \tag{A41}$$

Next, we can derive the relationship between $a$ and $\phi$ needed to achieve proofreading:

$$1 \geq \frac{(a\mu+m_s)\phi}{a(1-\mu+\phi\mu-\rho\phi)-(1-\phi)(w_{min}+m_s)} \tag{A42}$$

We can express this condition in terms of dimensionless quantities:

$$\frac{a}{w_{min}}(1-\mu+\phi\mu-\rho\phi)-(1-\phi)(1+\frac{m_s}{w_{min}}) \geq \left(\frac{a}{w_{min}}\mu+\frac{m_s}{w_{min}}\right)\phi \qquad \text{(A43)}$$

Solving for an exact condition on $\phi$, we find:

$$\frac{a}{w_{min}}(1-\mu-\rho\phi)-\left(1+\frac{m_s}{w_{min}}\right)+\phi \geq 0 \qquad \text{(A44)}$$

$$\frac{a}{w_{min}}(1-\mu)-\left(1+\frac{m_s}{w_{min}}\right) \geq \phi\left(\rho\frac{a}{w_{min}}-1\right) \qquad \text{(A45)}$$

$$\phi \leq \frac{\frac{a}{w_{min}}(1-\mu)-(1+\frac{m_s}{w_{min}})}{\frac{a}{w_{min}}\rho-1} \qquad \text{(A46)}$$

A few bounding relationships show that Condition A46 is more restrictive than A41:

$$\phi \leq \frac{\frac{a}{w_{min}}(1-\mu)-\left(1+\frac{m_s}{w_{min}}\right)}{\frac{a}{w_{min}}\rho-1} \leq \frac{\frac{a}{w_{min}}(1-\mu)-1}{\frac{a}{w_{min}}\rho-1} \leq \frac{\frac{a}{w_{min}}(1-\mu)}{\frac{a}{w_{min}}\rho} = \frac{1-\mu}{\rho} \qquad \text{(A47)}$$

Therefore, everywhere Condition A46 is satisfied, Condition A41 is also satisfied (and Condition A38 is satisfied). Condition A46 is all we need to draw the boundary between the two regimes, as shown in Figure 3 in the main text. We can re-arrange the same boundary to express it in terms of $a/w_{min}$:

$$\frac{a}{w_{min}} \geq \frac{1-\phi+\frac{m_s}{w_{min}}}{1-\mu-\rho\phi} \qquad \text{(A48)}$$

Lastly, we follow the same steps to find the boundary beyond which $x_1$ takes a certain value $x^*$.

$$\frac{a}{w_{min}} \geq \frac{\frac{m_s}{w_{min}}\phi+x^*(1-\phi)\left(1+\frac{m_s}{w_{min}}\right)}{x^*\left(1-\mu+\phi\mu-\frac{\mu\phi}{x^*}-\rho\phi\right)} \qquad \text{(A49)}$$

### B. Table of parameters tested in simulation

**Table S1.** Parameter values and ranges used for simulations.

| Parameter | Range | Rationale |
|---|---|---|
| $w_{min}$<br>Basal rate of cell death | 0.07 – 253 $y^{-1}$ | We estimated the range of $w_{min}$ from known turnover rates of cells and tissues. Cells in a tissue can have turnover rates as fast as 1–3 days (e.g., neutrophils, small intestine epithelial cells) or as slow as decades (e.g., cardiomyocytes).[1,2] Exceptions are lens cells, oocytes, and central nervous system neurons, which can last a lifetime. Turnover rate is defined as how long it takes to replace 50% of the cells. For our simulations, therefore, we test the range of turnover rate from 1 day to 10 years. This turnover rate translates to $w_{min}$ as follows: $$w_{min} = \frac{\ln(2)}{\text{turnover rate}}$$ |
| $a_0$<br>Proliferation rate of healthy cells | $w_{min}$ – $2w_{min}$ | From the model, we determine that the minimum $a_0$ to maintain tissue homeostasis is given by $a_0 = w_{min}$. We also find analytically that the boundary between model regimes depends on the ratio $a_0/w_{min}$. Therefore, in simulations, we vary $a_0$ as a multiple of $w_{min}$. Figure 3 shows the simulation using $a_0$ from $w_{min}$ to $2w_{min}$. |
| $\rho$<br>Neoplastic growth advantage | 1 – 2 | Prior histological and modeling work shows that neoplastic cells can divide up to twice as fast as healthy cells.[3–5] |
| $\phi$<br>False positive rate of apoptotic detector | 0 – 1 | Since $\phi$ is a theoretical parameter from our model, we consider every possible value of $\phi$ from zero to one. |
| $\mu$<br>The mutation rate per cell division | $10^{-7}$ – $10^{-4}$ | The range of mutation rates, from previous data.[5–8] Note that the $\mu$ we use in Figure 2 and Figure 3, $10^{-4}$, is on the high end of these estimates, in order to ensure that proofreading still works even under more conservative assumptions. |
| $m_s$<br>The rate of mutation occurring outside of cell division | 0 – $w_{min}$ | It is difficult to find estimates that disaggregate $\mu$ from $m_s$ in data. $m_s$ may be driven by environmental exposures, so we expect it to scale with $w_{min}$. Figure 3 shows how varying $m_s/w_{min}$ shifts the boundary between regimes without changing the steady state solutions in the high-flux limit (Equation A36). |

**C. Existence of non-zero steady states**

Tissue homeostasis requires that the population grows when $x_0 + x_1 < 1$, the population shrinks when $x_0 + x_1 > 1$, and the population stays the same size when $x_0 + x_1 = 1$. We start with the basic model (Equation A1-A2):

$$\frac{dx_0}{dt} = \boldsymbol{a_0} x_0 - \boldsymbol{w_0} x_0 - \boldsymbol{m} x_0 \tag{C1}$$

$$\frac{dx_1}{dt} = \boldsymbol{a_1} x_1 - \boldsymbol{w_0} x_1 + \boldsymbol{m} x_0 \tag{C2}$$

To assess the dynamics of the total population size, we consider the derivative of the sum with respect to time. The population stays the same size when this derivative is equal to zero:

$$\frac{d(x_0 + x_1)}{dt} = \boldsymbol{a_0} x_0 - \boldsymbol{w_0} x_0 + \boldsymbol{a_1} x_1 - \boldsymbol{w_1} x_1 \tag{C3}$$

We now define six functions: $\boldsymbol{f_{0+}}$ is a rate function associated with $x_0$ that is required to be zero unless $x_0+x_1 > 1;$ $\boldsymbol{f_{00}}$ is a rate function associated with $x_0$ that is required to be zero unless $x_0+x_1 = 1;$ and so on. Then tissue homeostasis requires:

$$0 = \boldsymbol{f_{00}}(x_0, x_1)x_0 + \boldsymbol{f_{10}}(x_0, x_1)x_1 \tag{C4}$$

$$0 > \boldsymbol{f_{0+}}(x_0, x_1)x_0 + \boldsymbol{f_{1+}}(x_0, x_1)x_1 \tag{C5}$$

$$0 < \boldsymbol{f_{0-}}(x_0, x_1)x_0 + \boldsymbol{f_{1-}}(x_0, x_1)x_1 \tag{C6}$$

These are the tissue homeostasis constraints summarized by Equation 7 in the main text. Conditions C5-C6 are necessary because Equation C4 alone does not guarantee the existence of a stable, nonzero steady-state population size. Equation C4 is satisfied if $x_0$ and $x_1$ are both zero. This is expected: in a population where cells only come from other cells, a population of zero will remain zero. Equation C4 is also satisfied if $\boldsymbol{f_{00}} = \boldsymbol{f_{10}} = 0$ for all $x$. However, because this condition is independent of $x$, the system in this scenario will have only marginal stability properties: if the total population size is perturbed, then it will retain its perturbed value, instead of tending back towards the homeostatic set point of population size. We must therefore verify that if Equation C4 is satisfied, the additional Conditions C5-C6 can also be satisfied.

This is straightforward to define piecewise because the functions are non-overlapping by definition. The simplest function that meets the requirements is a linear function of the vector $x$; this is one of many realistic compensation functions (i.e. continuous, simple, functions which can be implemented in molecular circuits). As long as the behavior on the line $x_0 + x_1 = 1$ is defined in such a way that it does not constrain the behavior off the line and vice versa, the behavior off the line can be defined in such a way that Conditions C5-C6 are met. Moreover, while we are analyzing the behavior on the line $x_0 + x_1 = 1$, as we do in our main analyses, Conditions C5-C6 do not affect the behavior. If there are steady states on the line $x_0 + x_1 = 1$, then the trajectory to the line does not affect the location or stability of those steady states. require feedback functions to simulate the model and to rule out trivial solutions to Equation C4. There are scenarios for which details of the feedback functions would affect the robustness of the homeostatic system as a whole, but for the scenarios analyzed in this study, any feedback function will serve the same role.

**D. Analysis for the case when a cell can accumulate multiple mutations**

We have defined $x_0$ as healthy cells and $x_1$ as cells with one significant mutation, in a framework that can extend out to $x_L$ as cells with $L$ significant mutations. In the main analysis, we work through the solutions for $L = 1$. In this section, we show how the key analytical results extend to $L > 1$. Increasing the number of mutations needed will make neoplasia rarer (as we confirm here), so these results are most useful to develop a richer quantitative sense of how the theoretical results might apply in real settings and how this additional suppression strategy interacts with those we highlight elsewhere. For some cases, such as the classical case of tumor suppression in pediatric retinoblastoma, evidence suggests a true biological $L$ of two.[7]

**Regime 1: Low-flux delaying**

We first derive the closed-form solution through time with $L>1$, with $a = a_0 = a_1\ldots$ and $w = w_0 = w_1\ldots$ along the hyperplane where the total population size is given by:

$$\sum_{l=0}^{L} x_l = 1 \tag{D1}$$

(D1 describes a hyperplane because it has $L$-$1$ degrees of freedom, defining a line in two dimensions, a plane in three dimensions, and so on.)

Starting from $L = 2$:

$$\frac{dx_0}{dt} = ax_0 - wx_0 - (a\mu + m_s)x_0 \tag{D2}$$

$$\frac{dx_1}{dt} = ax_1 - wx_1 + (a\mu + m_s)x_0 - (a\mu + m_s)x_1 \tag{D3}$$

$$\frac{dx_2}{dt} = ax_2 - wx_2 + (a\mu + m_s)x_1 \tag{D4}$$

If we sum across populations, we find:

$$\frac{d(x_0 + x_1 + x_2)}{dt} = a(x_0 + x_1 + x_2) - w(x_0 + x_1 + x_2) \tag{D5}$$

Tissue homeostasis requires that $x_0 + x_1 + x_2 = 1$ and that Equation D5 must be zero. Therefore, $a=w$ and the simplified dynamics are:

$$\frac{dx_0}{dt} = -(a\mu + m_s)x_0 \tag{D6}$$

$$\frac{dx_1}{dt} = (a\mu + m_s)x_0 - (a\mu + m_s)x_1 \tag{D7}$$

$$\frac{dx_2}{dt} = (a\mu + m_s)x_1 \tag{D8}$$

This description extends to arbitrarily many dimensions. Note that $x_L$ has slightly different dynamics from all of the preceding dimensions, so we use a compact matrix description for the preceding dimensions while leaving $x_L$ separate. Because the total population will sum to one, we can compute $x_L$ simply as $1-(x_0 + x_1 + ...)$.

$$\frac{dx_{0:L-1}}{dt} = Ax_{0:L-1} \tag{D9}$$

$$\frac{dx_L}{dt} = (a\mu + m_s)x_{L-1} \tag{D10}$$

$$A = \begin{bmatrix} -1 & 0 & 0 & 0 & 0 \\ 1 & -1 & 0 & 0 & 0 \\ 0 & 1 & \ddots & 0 & 0 \\ 0 & 0 & 1 & -1 & 0 \\ 0 & 0 & 0 & 1 & -1 \end{bmatrix}(a\mu + m_s) \tag{D11}$$

The solution to the problem described in Equations D9-D11, using matrix exponentiation, is:

$$x_L(t) = 1 - \mathrm{e}^{At}x_{0:L-1} \tag{D12}$$

$$x_{0:L-1}(0) = [1,0,0, \ldots 0] \tag{D13}$$

The repeating structure of $A$ allows a special solution to Equations D12-13. Specifically, to compute the total size across each subpopulation except $x_L$ at time $t$, we need to compute the sum of the first column of the exponentiated matrix. The matrix exponential is defined as:

$$e^{At} = I + At + \frac{(At)^2}{2!} + \frac{(At)^3}{3!} \ldots \tag{D14}$$

In the first column, the $\ell$-th row entry (indexed from zero) is given by:

$$e^{At}[0, l] = e^{-(a\mu+m_s)t}\frac{\left((a\mu + m_s)t\right)^{\ell}}{\ell!} \tag{D15}$$

$$x_L(t) = 1 - e^{-(a\mu+m_s)t}\sum_{l=0}^{L-1}\frac{\left((a\mu + m_s)t\right)^{\ell}}{\ell!} \tag{D16}$$

To interpret the indexing: $L = 2$ corresponds to three total states ($x_0, x_1, x_2$), so $A$ is a 2x2 or $LxL$ matrix. If $A$ is indexed from zero, then we sum from zero to $L-1$. As $L$ approaches infinity, the summation term becomes the Taylor expansion of the negative exponential, so $x_L$ approaches zero. It is readily verified that Equation D16 is identical to Equation A16 for $L = 1$.

**Regime 2: High-flux proofreading**

We now consider the case where a steady state exists for which $x_0$ is nonzero, with $L > 1$. In this case, we do not need to specifically consider linear coupling functions. We will assume that the division and death rates remain the same for all neoplastic cells. At steady state:

$$0 = a_0 x_0 - w_0 x_0 - m x_0 \quad \text{(D17)}$$
$$0 = a_1 x_1 - w_1 x_1 + m x_0 - m x_1 \quad \text{(D18)}$$
$$\ldots$$
$$0 = a_1 x_{L-1} - w_1 x_{L-1} + m x_{L-2} - m x_{L-1} \quad \text{(D19)}$$
$$0 = a_1 x_L - w_1 x_L + m x_{L-1} \quad \text{(D20)}$$

Therefore, because Equation D17 gives us that $a_0$-$w_0$=$m$, Equation D20 becomes:

$$0 = (a_1 - w_1) x_L + (a_0 - w_0) x_{L-1} \quad \text{(D21)}$$

To lighten notation, define:

$$W = \frac{a_0 - w_0}{w_1 - a_1} \quad \text{(D22)}$$

As long as a steady state exists, $W$ is guaranteed to be positive.

Substitute into Equation D21:

$$x_L = W x_{L-1} \quad \text{(D23)}$$

Moving back to Equation D19, divide through by $a_1$-$w_1$:

$$0 = x_{L-1} - W x_{L-2} + W x_{L-1} \quad \text{(D24)}$$
$$x_{L-1} = \frac{W}{1+W} x_{L-2} \quad \text{(D25)}$$

Noticing we can repeat the same relationship in Equations D24 and D25 for $x_{L-2}$ and $x_{L-3}$, we have the generic sequence up to $L$-$1$:

$$x_{k+1} = \frac{W}{1+W} x_k \quad \text{(D26)}$$

with the final step relating $x_L$ given by Equation D23. From Equation D18,

$$x_1 = \frac{W}{1+W} x_0 \quad \text{(D27)}$$

Putting these pieces together,

$$x_L = W\left(\frac{W}{1+W}\right)^{L-1} x_0 \tag{D28}$$

We now need the initial condition for this sequence, $x_0$. Using the fact that the total population should sum to 1:

$$1 = x_0 W\left(\frac{W}{1+W}\right)^{L-1} + x_0\sum_{\ell=0}^{L-1}\left(\frac{W}{1+W}\right)^{\ell} \tag{D29}$$

Because the summation term is a geometric series, we can make the following substitution:

$$\sum_{\ell=0}^{L-1}\left(\frac{W}{1+W}\right)^{\ell} = \frac{1-\left(\frac{W}{W+1}\right)^{L}}{1-\frac{W}{W+1}} \tag{D30}$$

Therefore:

$$\frac{1}{x_0} = W\left(\frac{W}{1+W}\right)^{L-1} + \frac{1-\left(\frac{W}{W+1}\right)^{L}}{1-\frac{W}{W+1}} \tag{D31}$$

$$\frac{1}{x_0} = W\left(\frac{W}{1+W}\right)^{L-1} + W + 1 - (W+1)\left(\frac{W}{W+1}\right)^{L} \tag{D32}$$

$$\frac{1}{x_0} = W\left(\frac{W}{1+W}\right)^{L-1} + W + 1 - (W+1)\frac{W}{W+1}\left(\frac{W}{W+1}\right)^{L-1} \tag{D33}$$

$$\frac{1}{x_0} = W + 1 \tag{D34}$$

$$x_0 = \frac{1}{W+1} \tag{D35}$$

Recall that our ultimate goal is to find $x_L$, which we previously found to be:

$$x_L = W\left(\frac{W}{1+W}\right)^{L-1} x_0 \tag{D36}$$

Therefore,

$$x_L = \frac{W}{1+W}\left(\frac{W}{1+W}\right)^{L-1} \tag{D37}$$

$$x_L = \left(\frac{W}{1+W}\right)^{L} \tag{D38}$$

Substituting back in for *W,*

$$x_L = \left( \frac{\frac{a_0 - w_0}{w_1 - a_1}}{1 + \frac{a_0 - w_0}{w_1 - a_1}} \right)^L \tag{D39}$$

$$x_L = \left( \frac{a_0 - w_0}{w_1 - a_1 + a_0 - w_0} \right)^L \tag{D40}$$

As a common-sense check, Equation D40 reduces to Equation A24 when $L = 1$.

**E. Generalizing the model analysis to the case of nonlinear couplings**

At steady state, the rate functions are constant, and the following equations hold (Equations A1-A2 at steady state):

$$0 = a_0 x_0 - w_0 x_0 - m x_0 \tag{E1}$$
$$0 = a_1 x_1 - w_1 x_1 + m x_0 \tag{E2}$$

Summing Equations E1 and E2, we find:

$$0 = a_0 x_0 - w_0 x_0 + a_1 x_1 - w_1 x_1 \tag{E3}$$
$$w_1 x_1 - a x_1 = a_0 x_0 - w_0 x_0 \tag{E4}$$

Rearranging Equations E3–E4 to solve for the ratio $x_1/x_0$, in the scenario where at least one non-zero steady-state $x_0$ exists,

$$\frac{x_1}{x_0} = \frac{a_0 - w_0}{w_1 - a_1} \tag{E5}$$

Substituting Equation E1 into Equation E5,

$$\frac{x_1}{x_0} = \frac{m}{w_1 - a_1} \tag{E5}$$

To confirm that flux-based proofreading is possible without the assumption of linear coupling, we should confirm that $x_1$ is made small when $a_0$ is small (i.e., increasing proliferation rate suppresses neoplastic cells). We can solve for the steady state directly, using Equation E2 and the additional constraint that $x_0 + x_1 = 1$:

$$0 = a_1 x_1 - w_1 x_1 - m(1 - x_1) \tag{E7}$$
$$x_1 = \frac{m}{w_1 - a_1 + m} \tag{E8}$$

Along the line $x_0 + x_1 = 1$ (where Equation E8 holds), if $x_1$ is made small with increases in $a_0$, then the ratio $x_1/x_0$ is also made small with increases in $a_0$. Likewise, if $x_1$ approaches zero, $x_1/x_0$ approaches zero. This confirms that it is sufficient for our purposes to consider Equation E6, which is simpler and more interpretable.

To determine where increasing $a_0$ decreases $x_1$, we can take the partial derivative in Equation E6 of $x_1/x_0$ with respect to $a_0$. We use prime notation to represent this partial derivative:

$$\left(\frac{x_1}{x_0}\right)' = \frac{(w_1 - a_1)m' - m(w_1 - a_1)'}{(w_1 - a_1)^2} \tag{E9}$$

We see from Equation E9 that if the derivative in left-hand side is negative, then increasing $a_0$ reduces $x_1/x_0$. Because the denominator of the right-hand side in Equation E9 is the square of a real quantity (and therefore always positive), the condition for which increasing $a_0$ suppresses neoplasia is satisfied when the numerator takes negative values:

$$(w_1 - a_1)m' < m(w_1 - a_1)' \quad \text{(E10)}$$

$$\frac{m'}{m} < \frac{(w_1 - a_1)'}{(w_1 - a_1)} \quad \text{(E11)}$$

Anywhere Condition E11 is satisfied, increasing flux will increase proofreading, decreasing the steady-state fraction of neoplastic cells. For a given mutation-proliferation coupling, what needs to be true about the discrimination-proliferation coupling to guarantee that $x_1$ is small in the high proliferation limit? The general principle is illustrated by polynomial functions, or functions that are well-approximated by polynomial functions: in the limit as $a_0$ goes to infinity, the leading term of the polynomial will dominate. Therefore, if both the mutation-proliferation coupling and the discrimination-proliferation coupling are both order $q$ with coefficients $c$ and $d$ respectively,

$$m = \sum_{i=0}^{q} c_i a_0^i \quad \text{(E12)}$$

$$w_1 - a_1 = \sum_{i=0}^{q} d_i a_0^i \quad \text{(E12)}$$

then the ratio $x_1/x_0$ will be $c_q/d_q$, and small steady state $x_1$ will depend on how much larger $d_q$ is than $c_q$. This is the scenario in the linear coupling case. If the discrimination-proliferation coupling is higher-order than the mutation-proliferation coupling (e.g. linear mutation to quadratic discrimination, or an exponential to any polynomial), the ratio $x_1/x_0$ will be zero and steady state $x_1$ will go to zero with large $a_0$.

In general, increasing proliferation $a_0$ will increase the mutation rate $m$. Proliferation suppresses neoplasia when the net death rate of neoplastic cells ($w_1$-$a_1$) increases faster than the mutation rate, and completely suppresses neoplasia if net death rate dominates the mutation rate in the high-proliferation limit. In the proofreading regime of the linear case, the net death rate grows faster, but does not dominate the mutation rate. In the nonlinear case, the net death rate can theoretically dominate the mutation rate, driving $x_1$ to zero.

We expect these coupling functions to eventually saturate in real systems: that is, additional physiological constraints on rates will mean that increasing proliferation after some point will have diminishing returns on discrimination. Therefore, the useful interpretation of Equation E6 and Condition E11 is for local changes: Will increasing proliferation from its current level improve discrimination? Is the current rate of discrimination much larger than the current rate of mutation? These questions will determine the system behavior.

**F. The model with stem vs differentiated cells**

In the main model, all cells divide and die. This simplification emphasizes the key principles. However, in most real cell populations, some cells are typically terminally differentiated and do not divide, carrying out other physiological functions. (There are some important exceptions, like immune cell populations.) In this section, we consider the effects of modeling differentiated (non-dividing) vs stem (dividing) cell populations. We model a differentiated population $x_{d0}$ that is supplied by the stem population $x_{s0}$ with a differentiation rate $\boldsymbol{r}$. A neoplastic population $x_{d1}$ is similarly supplied by the population $x_{s1}$, but this population retains a self-renewing capacity. This models the common biological scenario where stem cells are compartmentally separated from differentiated cells. The tissue homeostasis constraint should now require $x_{d0} + x_{d1} = 1$, since this is the functional compartment. Spontaneous mutation through environmental exposures can transform cells from $x_{d0}$ to $x_{d1}$, but this rate may differ between the stem cell compartment and the differentiated compartment With these definitions, we write the following model:

$$\frac{dx_{s0}}{dt} = \boldsymbol{a_0} x_{s0} - \boldsymbol{w_0} x_{s0} - \boldsymbol{r} x_{s0} - \boldsymbol{m} x_{s0} \tag{F1}$$

$$\frac{dx_{d0}}{dt} = \boldsymbol{r} x_{s0} - \boldsymbol{w_{d0}} x_{d0} - m_{ds} x_{d0} \tag{F2}$$

$$\frac{dx_{s1}}{dt} = \boldsymbol{a_1} x_{s1} - \boldsymbol{w_1} x_{s1} + \boldsymbol{m} x_{s0} - \boldsymbol{r} x_{s1} \tag{F3}$$

$$\frac{dx_{d1}}{dt} = \boldsymbol{a_{d1}} x_{d1} - \boldsymbol{w_{d1}} x_{d1} + \boldsymbol{r} x_{s1} + m_{ds} x_{d0} \tag{F4}$$

To consider whether flux-based proofreading consider the steady state. As in Section E, we can solve for steady-state $x_{d1}/x_{d0}$ as a way to find steady-state $x_{d1}$.

$$0 = a x_{s0} - w_0 x_{s0} - r x_{s0} - m x_{s0} \tag{F5}$$

$$0 = r x_{s0} - w_{d0} x_{d0} - m_{ds} x_{d0} \tag{F6}$$

$$0 = a_1 x_{s1} - w_1 x_{s1} + m x_{s0} - r x_{s1} \tag{F7}$$

$$0 = a_{d1} x_{d1} - w_{d1} x_{d1} + r x_{s1} + m_{ds} x_{d0} \tag{F8}$$

Rearranging Equations F6 and F7, we obtain the following expressions:

$$\frac{x_{d0}}{x_{s0}} = \frac{r}{w_{d0} + m_{ds}} \tag{F9}$$

$$\frac{x_{s1}}{x_{s0}} = \frac{m}{r + w_1 - a_1} \tag{F10}$$

Then, dividing Equation F8 through by $x_{d0}$,

$$0 = (a_{d1} - w_{d1}) \frac{x_{d1}}{x_{d0}} + r \frac{x_{s1}}{x_{d0}} + m_{ds} \tag{F11}$$

$$\frac{x_{d1}}{x_{d0}} = \frac{1}{(w_{d1} - a_{d1})} \left(r \frac{x_{s1}}{x_{d0}} + m_{ds}\right) \tag{F12}$$

We use Equations F9 and F10 to expand Equation F12.

$$\frac{x_{d1}}{x_{d0}} = \frac{1}{(w_{d1} - a_{d1})}\left(r\left(\frac{m}{r + w_1 - a_1}\right)\left(\frac{w_{d0} + m_{ds}}{r}\right) + m_{ds}\right) \tag{F13}$$

Simplifying Equation F13,

$$\frac{x_{d1}}{x_{d0}} = \frac{1}{(w_{d1} - a_{d1})}\left(\frac{m(w_{d0} + m_{ds})}{r + w_1 - a_1} + m_{ds}\right) \tag{F14}$$

Now, we substitute in the linear coupling functions.

$$\frac{x_{d1}}{x_{d0}} = \frac{1}{(w_{min} + k - a\rho)}\left(\frac{(a\mu + m_s)(w_{min} + k\phi + m_{ds})}{r + w_{min} + k - a\rho} + m_{ds}\right) \tag{F15}$$

This leaves $k$ as the last $a$-dependent term that is not explicitly accounted for. To solve for $k$, we use Equation F5:

$$k = \frac{a(1 - \mu) - w_{min} - r - m_s}{\phi} \tag{F16}$$

The model described by Equations F15 and F16 has more degrees of freedom than the model in the main text. Fully exploring this model's behaviors is beyond the scope of this simple confirmatory analysis. As before, within the relevant parameter regime, $x_{d1}$ approaches a lower bound as $a$ increases. Therefore, high-flux proofreading is possible within this model. To further characterize high-flux proofreading in this model, we take the limit of Equation F16:

$$\lim_{a\to\infty} \frac{x_{d1}}{x_{d0}} = \frac{1}{\left(\frac{a(1-\mu)}{\phi} - a\rho\right)}\left(\frac{a\mu\left(\frac{a(1-\mu)}{\phi}\phi\right)}{\frac{a(1-\mu)}{\phi} - a\rho}\right) \tag{F17}$$

Then, simplifying, we find:

$$\lim_{a\to\infty} \frac{x_{d1}}{x_{d0}} = \frac{\mu(1-\mu)\phi^2}{(1 - \mu - \rho\phi)^2} \tag{F18}$$

We solve for $x_{d1}$ using $x_{d0} + x_{d1} = 1$.

$$\lim_{a\to\infty} x_{d1} = \frac{\mu(1-\mu)\phi^2}{(1 - \mu - \rho\phi)^2 + \mu(1-\mu)\phi^2} \tag{F19}$$

Compare this to Equation A36, rewritten here:

$$\lim_{a\to\infty} x_1 = \frac{\mu\phi}{1-\mu+\phi\mu-\rho\phi} \tag{F20}$$

We can develop additional intuition with the approximation $1\text{-}\mu \approx 1$. Because we are interested in the case where $\rho \geq 1$, we use the same reasoning to approximate $\rho\text{-}\mu \approx \rho$.

$$\lim_{a\to\infty} x_1 \approx \mu\frac{\phi}{1-\rho\phi+\mu\phi} \tag{F21}$$

$$\lim_{a\to\infty} x_{d1} \approx \frac{\mu\phi^2}{(1-\rho\phi)^2+\mu\phi^2} \tag{F22}$$

Now suppose that $\rho\phi$ is small enough that $(1\text{-}\ \rho\phi)^2$ is suitably larger than $\mu$, another assumption that is well-supported in the numerical ranges we consider. In that case:

$$\lim_{a\to\infty} x_1 \approx \mu\frac{\phi}{1-\rho\phi} \tag{F23}$$

$$\lim_{a\to\infty} x_{d1} \approx \mu\left(\frac{\phi}{1-\rho\phi}\right)^2 \tag{F24}$$

These approximate bounds now have a parallel structure, with the additional steps of the stem cell model reflected by the square in Equation F24. This square will either increase or decrease the bound depending on the particular $\rho$ and $\phi$, and parameters that make $x_{d1}$ smaller are well within the range we consider. Therefore, high-flux proofreading is possible in a model with stem cells, and for some parameters, the model with stem cells can also improve proofreading performance.
